# Inter-individual differences in cerebrovascular reactivity are synchronized within functional networks and tissue layers: evidence from healthy older adults and patients with hypertension

**DOI:** 10.1101/2024.06.09.598005

**Authors:** Nuwan D. Nanayakkara, Liesel-Ann Meusel, Nicole D. Anderson, J. Jean Chen

**Affiliations:** Rotman Research Institute, Baycrest Academy for Research and Education; Departments of Psychology and Psychiatry, University of Toronto; Department of Medical Biophysics, University of Toronto; Department of Biomedical Engineering, University of Toronto

## Abstract

Functional-connectivity mapping has primarily relied resting-state functional MRI (rs-fMRI), and resting-state functional networks (RSNs) have been used widely to represent interactions within brain circuits. However, recent work demonstrated that resting-state functional networks (RSNs) may co-exist with vascular networks. In this work, we clarify the nature of these vascular networks by assessing the spatial covariation structure in breath-holding-based CVR amplitude and lag in a group of healthy older adults. We demonstrate a spatial synchrony in CVR amplitude and lag co-variations across participants confined to RSNs. Such a network structure is not seen when looking at the time-variate BOLD signal response to the breathhold. a network structure is also maintained in older adults with clinical hypertension, demonstrating its robustness against vascular pathologies. CVR amplitude is also found to vary with tissue layer in the grey matter and white matter, being most variable in deep WM and least variable in superficial cortex. Conversely, CVR lag appears to be organized by fibre tracts. This work demonstrates the use of cross-participant covariation patterns in CVR as an informative way of mapping the vascular routes in the GM and WM, and also raises questions about the nature and interpretation of RSNs.

## Introduction

The discovery of resting-state fMRI connectivity (rs-fcMRI) has led to the rapid development of brain-network (RSN) mapping, whereby spatial covariations in the BOLD fMRI signal are taken as indication of neuronal connections. While the existence of subcortical white-matter connections have provided strong support for the neuronal contribution to these networks, there is accumulating evidence that these network structures are not exclusively neuronal.

Bright et al. (Bright et al., 2020) used independent component analysis (ICA) to decompose the BOLD fMRI signal during simultaneous functional and breath-hold tasks and found that both decompositions consisted of overlapping networks that responded to two different stimuli, respectively. Tong et al. (Tong and Frederick, 2010) demonstrated the propagation of peripheral (i.e. systemic) vascular oscillations through the whole brain by way of the resting-state BOLD fMRI signal. They further demonstrated that systemic vascular oscillations that incorporate region-specific time-shifts alone (lag map) will result in patterns of ICA-derived spatial synchrony similar to the RSNs, irrespective of the underlying neuronal activity (Tong et al., 2015). Using similar methods and more specifically focusing on the contribution of low-frequency physiological fluctuations (i.e. cardiac and respiratory fluctuations), Chen et al. demonstrated the existence of robust physiological networks that resemble the widely published neuronal networks (Chen et al., 2020). Interestingly, it was found that the “physiological” signal fluctuations that give rise to these networks are spatially heterogeneous and thus cannot be removed by single-regressor methods. These works echo findings by Bright et al. who surmised that by coordinating blood flow across brain regions that typically exhibit synchronous neuronal activity, such vascular networks would also provide the most efficient hemodynamic support for increased network metabolism. Thus, the evidence suggests that such coordination is also observable during a breath-hold task.

The breath-hold BOLD response is essentially a measure of cerebrovascular reactivity (CVR). CVR is a spatially heterogeneous property of cerebral vasculature that can govern the amplitude and latency in task-based and resting-state BOLD responses (Chen and Gauthier, 2021). It can also govern the extent of the physiological manifestation in the BOLD signal (Chu et al., 2018; Golestani et al., 2016; Liu et al., 2013). As CVR can be further described by its magnitude and time lag, these CVR metrics can allow us to further dissect the nature of these network-like “vascular” and “physiological” synchronies. However, it is unclear whether such network structures exist in the fMRI time series corresponding to a vascular challenge, or in the CVR measurements themselves. It is also unclear whether vascular pathologies such as hypertension affect the spatial distribution and structure of CVR amplitude and lag. The goal of this work is two-fold: (1) to illustrate the existence (or absence) of network structure in CVR metrics in healthy older adults; (2) to determine whether vascular pathology such as found in hypertension affects the network structure in CVR variations across individuals.

## Methods

### Participants

Older adults of ages 65-85 were recruited and placed in control (CTL) and hypertension (HT) groups based on screening measures. The study was approved by the institutional Research Ethics Board. Participants were excluded from the study if they met any of the following criteria: (1) a score ≤ 31 on the Telephone Interview for Cognitive Status – modified version (Welsh et al., 1993) in order to exclude participants with possible dementia; (2) the presence of diabetes mellitus, based on self-report, including clinically significant gastroparesis, retinopathy, nephropathy, neuropathy, hepatic disease, or a recent coronary heart disease event as determined by a physician; (3) other significant medical or psychiatric disorders affecting cognitive function, such as stroke and major depressive disorder; (4) current or recent use of central nervous system-active medications, including those for the treatment of depression, sleep disorders, and migraine headaches; (5) major inflammatory disorders, heart failure, and chronic lung disease; or (6) hormone replacement therapy in female participants. The included participants were screened to ensure group status as listed below:

- CTL: Participants had a mean systolic BP ≤ 140 mmHg, a mean diastolic BP ≤ 90 mmHg, no history of antihypertensive medication use, and a fasting glucose level (FGL) ≤ 6.1 mmol/L.
- HT: Participants were using antihypertensive medication under physician prescription for a minimum of two years, with current blood pressure within a normal or HT range and limited to those who were using long-acting antihypertensive medications (e.g., ACEIs, angiotensin II receptor blockers, diuretics) in order to capture the most commonly prescribed medications.

Available data from 21 CTL:21 and 23 HT participants were preprocessed. Data from 3 CTLs and 3 HTs were removed from analysis due to excessive head-motion artifacts larger than 1^0^ rotation and 1 mm translation as detected by FSL motion correction. This study consisted of remaining data from 18 CTLs (M/F ratio = 1.25, age = 70.2 ± 3.3 years) and 20 HTs (M/F ratio = 0.43, age = 71.9 ± 4.7).

### Data acquisition

Each participant followed a set of 6 repetitions of a 30 s resting and 2 s exhale followed by a 15 s BH guided by visual clues (total duration, *T* = 47 s) during a dual-echo pCASL fMRI image acquisition session on a Siemens Trio 3T system (T2*-weighted echo-planar imaging, FOV = 220 mm, acquisition matrix = 220 x 220, voxel size = 3.4 x 3.4 x 6.0 mm, bandwidth = 2790 Hz/Pixel, TE1/TE2/TR = 9.1/25/4000 ms, flip angle = 90 degrees, slices = 16, averages = 1, concatenations = 1, scan duration = 5:24). The data associated with the second TE were used to compute the BOLD time series. Respiratory bellows were recorded using the scanner’s built-in belt. A T1 anatomical scan (FOV = 256 mm, acquisition matrix = 192 x 256, voxel size = 1.0 mm^3^, bandwidth = 200 Hz/Pixel, TI/TE/TR = 1100/2.63/2000 ms, flip angle = 9 degrees, slices = 160) was acquired for anatomical reference and tissue segmentation.

### CVR mapping

The dual-echo pCASL time series data were preprocessed using FSL (Jenkinson et al., 2012; Smith et al., 2004) and AFNI (Cox, 1996; Cox and Hyde, 1997) tools, and steps include slice timing correction, motion correction (for the BOLD echo separately) and registration into MNI space using *flirt*. The BOLD time series was obtained by surround averaging to produce 78 frames of BOLD data, which were then high-pass filtered to remove low-frequency noise from the data with a 0.01 Hz cut-off frequency, resulting in the “preprocessed BOLD signal”.

CVR amplitude and lag were estimated using a Fourier-based method we previously published (Nanayakkara et al., 2023). Briefly, CVR amplitude is identified from the Fourier spectrum of the BOLD time series during the BH task, and CVR lag is determined as the phase of the Fourier spectrum referenced to the minimal phase in the whole-brain volume, which is in turn computed using the phase of the global-mean signal as reference. For more details please refer to (Nanayakkara et al., 2023).

All CVR maps and time-series BOLD data were then transformed into MNI152 space using non-rigid registration (FSL *fnirt*) for the group statistical analysis.

### Statistical analysis

#### Time-series analysis

##### Region-of-interest-based correlation analysis

We segregated the data into regions of interest (ROIs) based on the Functionnectome (Nozais et al., 2021), which provides the masks for 30 functional networks in the grey matter (GM) and the corresponding white-matter (WM) voxels informed by the cortical network localizations and underlying tractography. Thus, each GM ROI has a corresponding WM ROI. Some sample ROIs are shown in **Fig. 1**, and the names of all 30 networks are summarized in **Table 2**. To minimize partial-volume effects between GM and WM in these ROIs, we isolated the GM by binarizing it in the MNI152 parcellations, and then subtracted the whole-brain mask by the cerebrospinal fluid mask and a dilated GM mask (by 2 mm) to obtain the WM mask. This approach prevented the WM mask from being over eroded. We then applied these masks to the Funnectome GM and WM ROIs; all Functionnectome network ROIs were thresholded at z > 7. Furthermore, to test whether each network ROI is in fact the best way to define a specific CVR pattern, we further subdivided each network ROI by randomly sampling voxels into 10 non-overlapping sub-ROIs, which would allow us to detect strong cross-network associations in the BOLD time series, should it exist. The time series in each of these 300 ROIs were averaged across voxels, and concatenated across all CTL participants to compute Pearson’s correlation matrices, as summarized in **Fig. 3**.

**Table 1.** Participant demographics.

| Demographic Variable | Control (CTL) group<br>(n = 18) | Hypertension (HT) group<br>(n = 20) |
| --- | --- | --- |
| Age (years) | $70.2 \pm 3.3$ | $71.9 \pm 4.7$ |
| Male/female ratio | 1.25 | 0.43 |
| Systolic BP (mmHg) | $123.2 \pm 13.1$ | $133.9 \pm 13.6$ |
| Diastolic BP (mmHg) | $74.1 \pm 5.8$ | $78.5 \pm 9.8$ |
| Hypertension Duration (years) | – | $10.6 \pm 8.7$ |

**Table 2.**
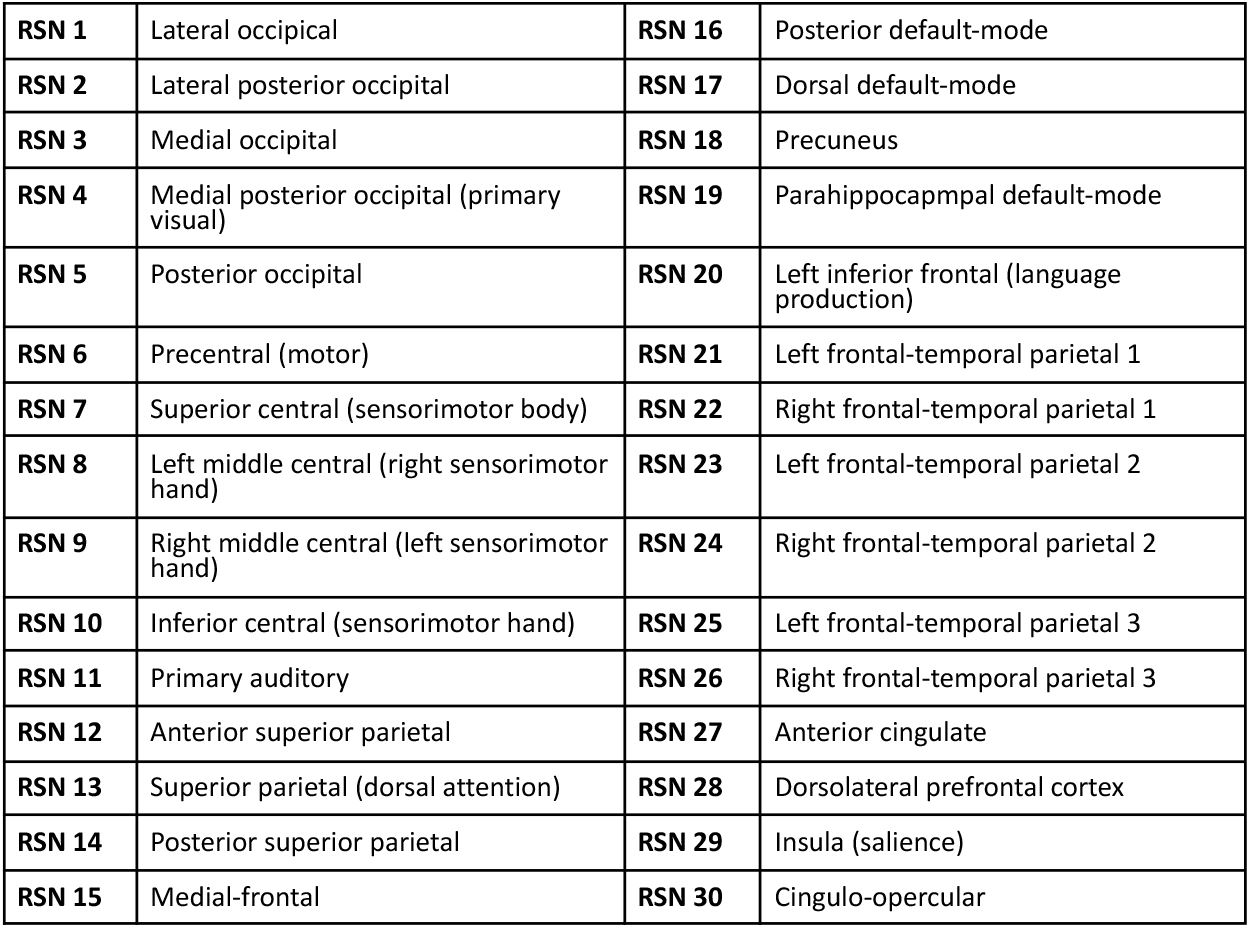
List of network ROIs of the Funnectionnectome.

**Figure 1.**
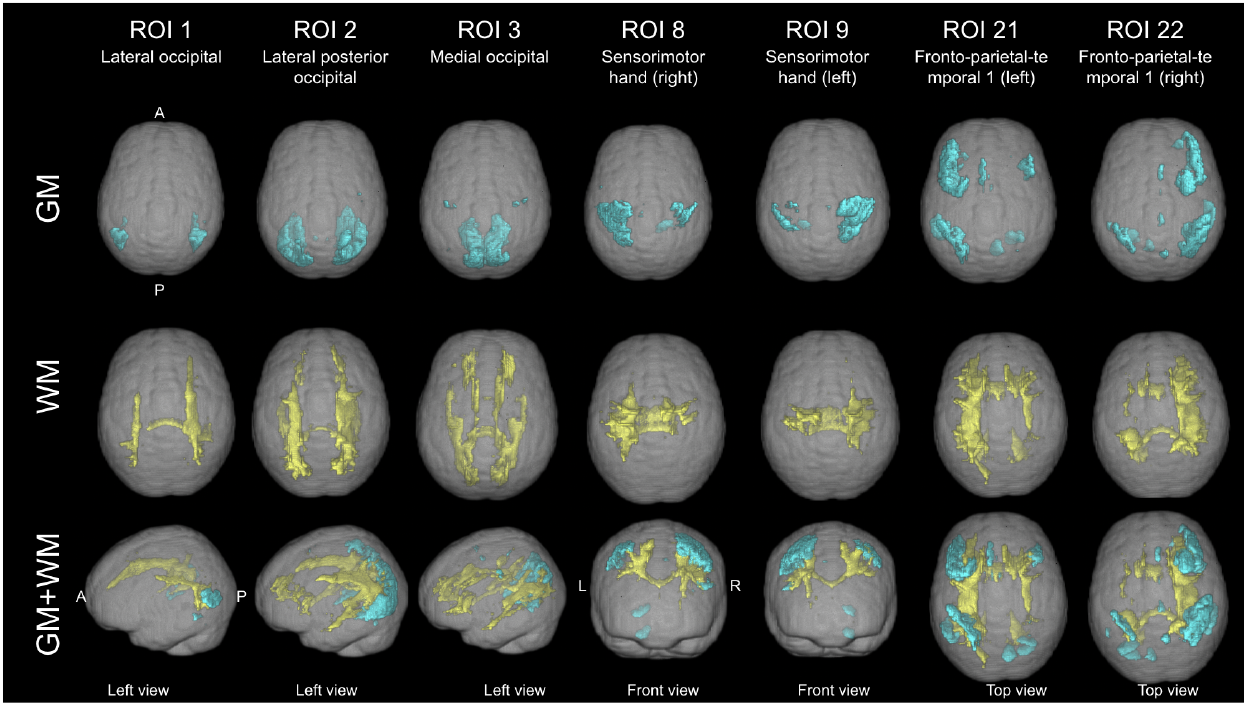
3D renderings of sample GM (blue) and corresponding WM (yellow) networks in the Functionnectome. These views confirm that the overlap between the GM and WM is negligible.

**Figure 2.**
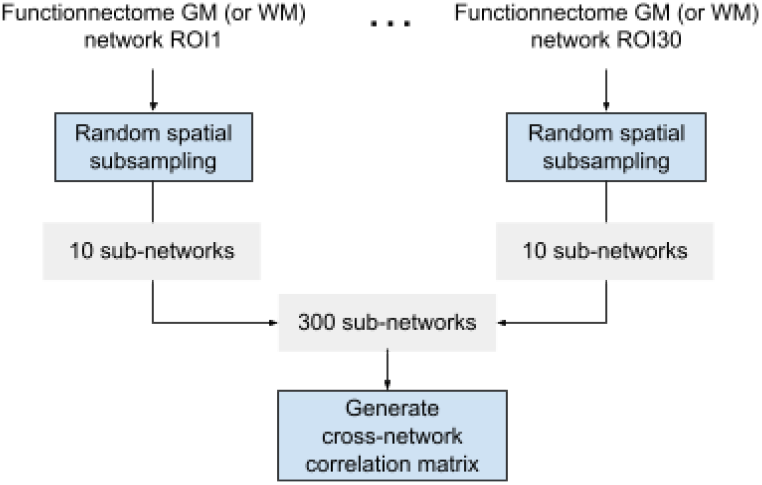
Flow chart for ROI generation.

**Figure 3.**
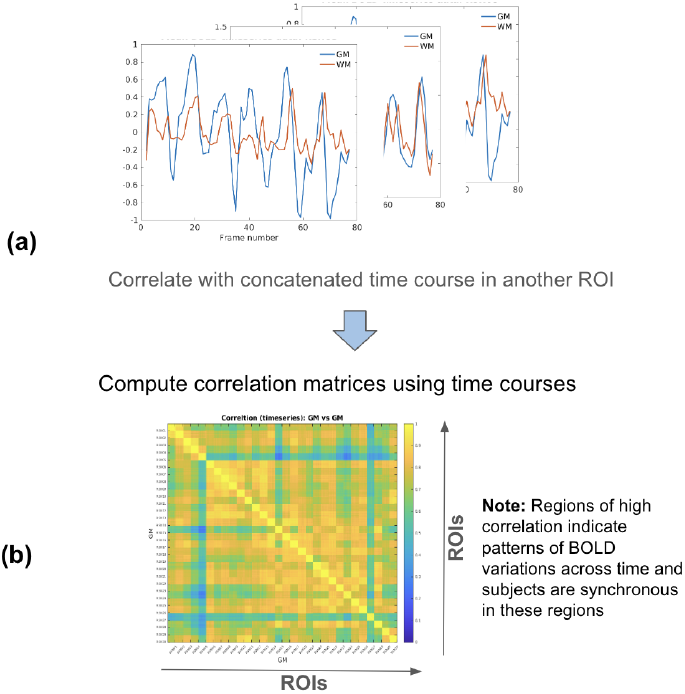
Flow chart for BH time-series correlation matrices. The time series are retracted from the ROIs defined in **Fig. 2**.

##### independent component analysis

As an initial approach, we performed group-based independent component analysis (ICA) on data from the CTL group. This was performed using FSL’s *melodic* (FSL Version 6.0.6) (Jenkinson et al., 2012). The resultant ICs were thresholded at *p < 0*.*01*, and visually inspected.

#### Cross-participant analyses

##### Region-of-interest-based correlation analysis

The CVR amplitude and lag of each participant was identified as the voxel-wise average value in each of the ROIs described earlier. In each of the 300 ROIs, the CVR amplitude and lag variations across participants were averaged across voxels, to enable the computation of cross-participant correlation matrices in the CTL and HT groups separately, as summarized in **Fig. 4**. The difference between the CTL and HT-based correlation matrices was assessed using the z test.

**Figure 4.**
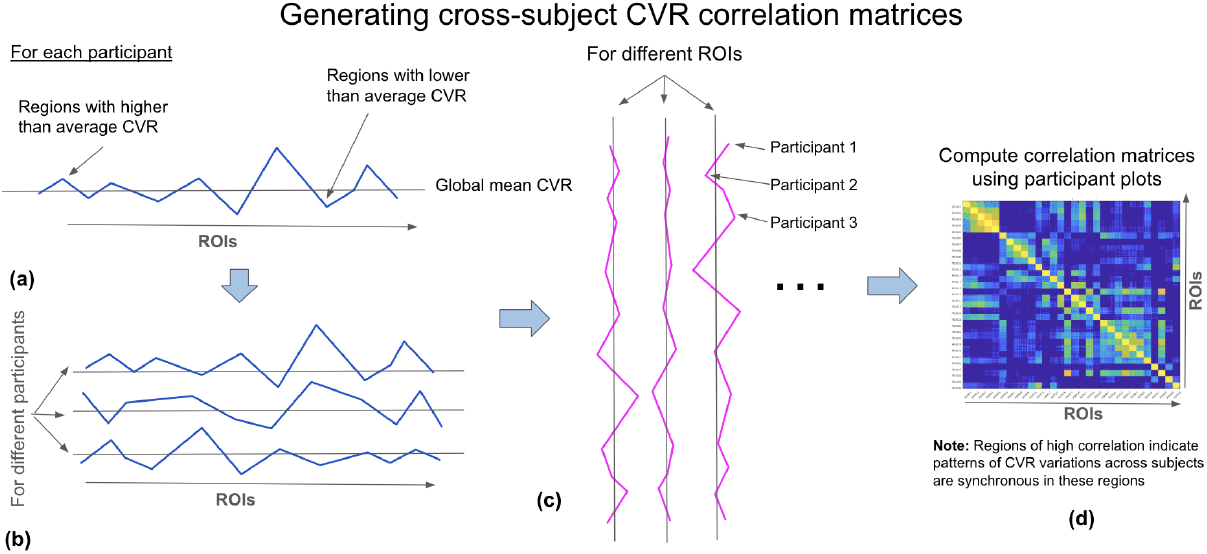
Flow chart for cross-participant CVR correlation matrices. The cross-participant CVR variations are retracted from the ROIs defined in **Fig. 2**.

##### Voxel-wise cluster analysis

The CVR amplitude and lag maps from all CTL participants were concatenated into a 4D volume, on which k-means clustering was performed based on Euclidean distance, implemented in Matlab (Natick, USA). We used k of 7 and 17 clusters, inspired by the numbers of networks in the Yeo atlas (Yeo et al., 2011), as well as k of 30, consistent with the Functionnectome. The clustering was also performed on 4D volumes consisting of GM and WM separately.

## Results

The ICs extracted from the concatenated BH time series reveal several interesting patterns (**Fig. 5**). For instance, In the GM, we extracted ICs that cover the entire cortex (**Fig. 5a, b**), the medial visual network (**Fig. 5c**), the lateral visual network (**Fig. 5d**), the basal ganglia network (**Fig. 5e, f**). In the WM, the ICA uncovered the left and right WM underlying the frontal attention network (**Fig. 5g, h**), the posterior thalamic radiations (**Fig. 5j**), the WM underlying the frontal-parietal network (**Fig. 5k**), the superior temporal WM (**Fig. 5l**) and the anterior corona radiata (**Fig. 5m**). A spatial overlap analysis was conducted between all ICs and the Funnctionnectome atlas, and the results shown in **Fig. 5n,o**, demonstrating that while there was more overlap between the WM ICs and WM networks than between GM ICs and GM networks.

**Figure 5.**
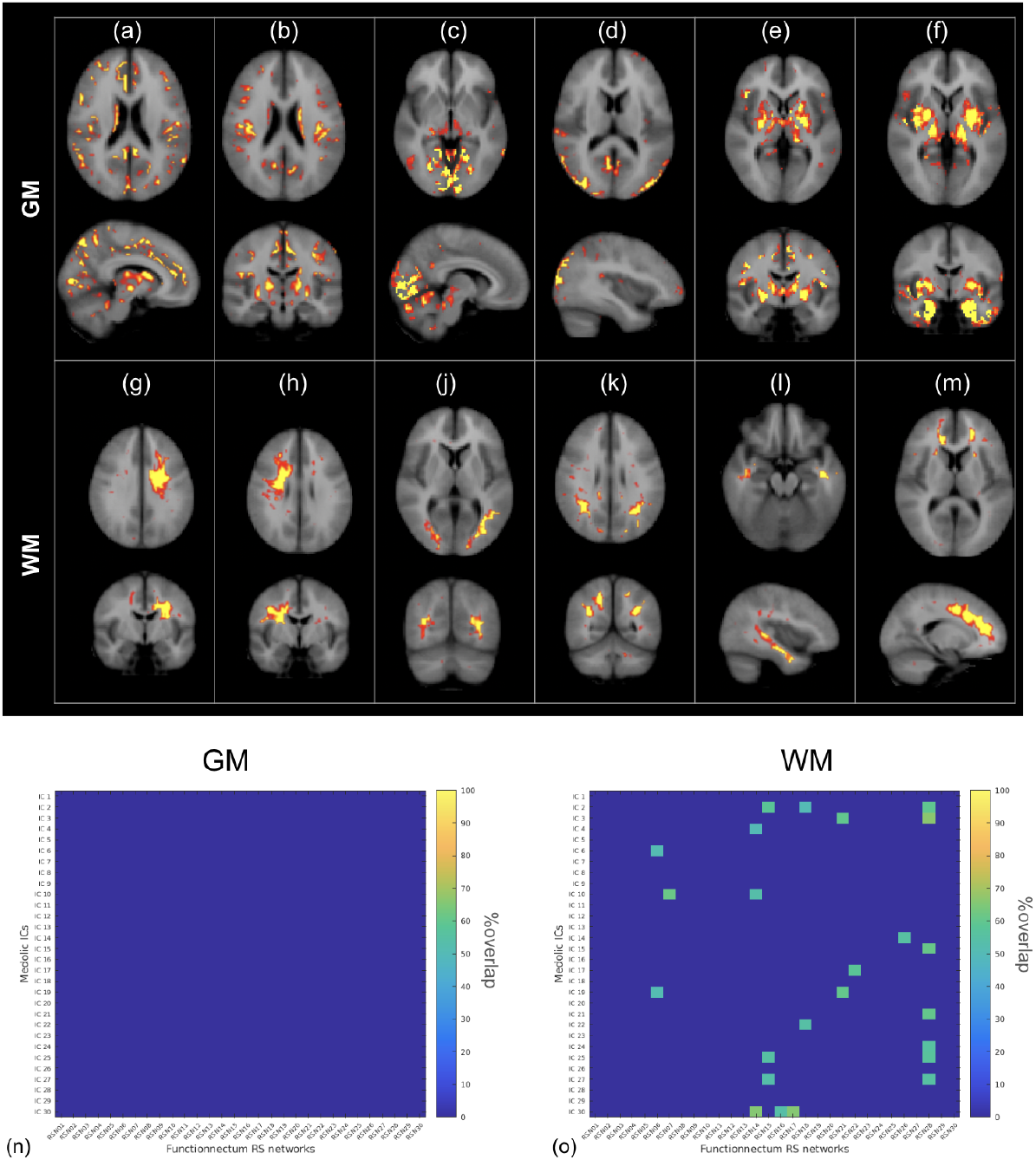
Sample ICs from the group ICA on BOLD time series data in the GM (a-f) and WM (g-m) in CTL datasets. The spatial overlaps between all 30 ICs and the 30 Funnectome network ROIs in GM (n) and WM (o) are also shown. The percentage of overlaps are thresholded at 50%.

In terms of BOLD time courses during the BH challenge (**Fig. 6**), most ROIs across the GM and nearly all ROIs in the WM appear to be uniformly correlated with each other. Nonetheless, in the GM, the correlation values are highest in the diagonal, suggesting that BH-related BOLD time courses are most synchronous within the same networks.

**Figure 6.**
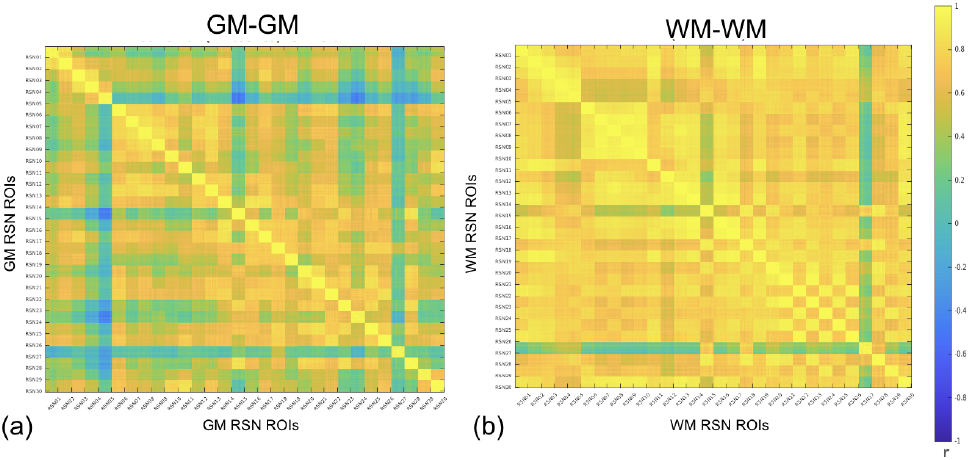
BOLD time series correlation matrix for CTL. (a) GM, (b) WM. Each grid point consists of 10 sub-grid points, representing the 10 randomly subsampled ROIs for each RSN.

Unlike in the case of the BOLD time series, the CVR metrics demonstrate distinctly network-specificity in the GM, in that CVR variations across participants are most synchronous across ROIs that belong to the same network (in the diagonal squares) (**Fig. 7**). GM CVR amplitude and lag correlations display very similar spatial structures. This remains the case for the WM, but to a lesser extent.

**Figure 7.**
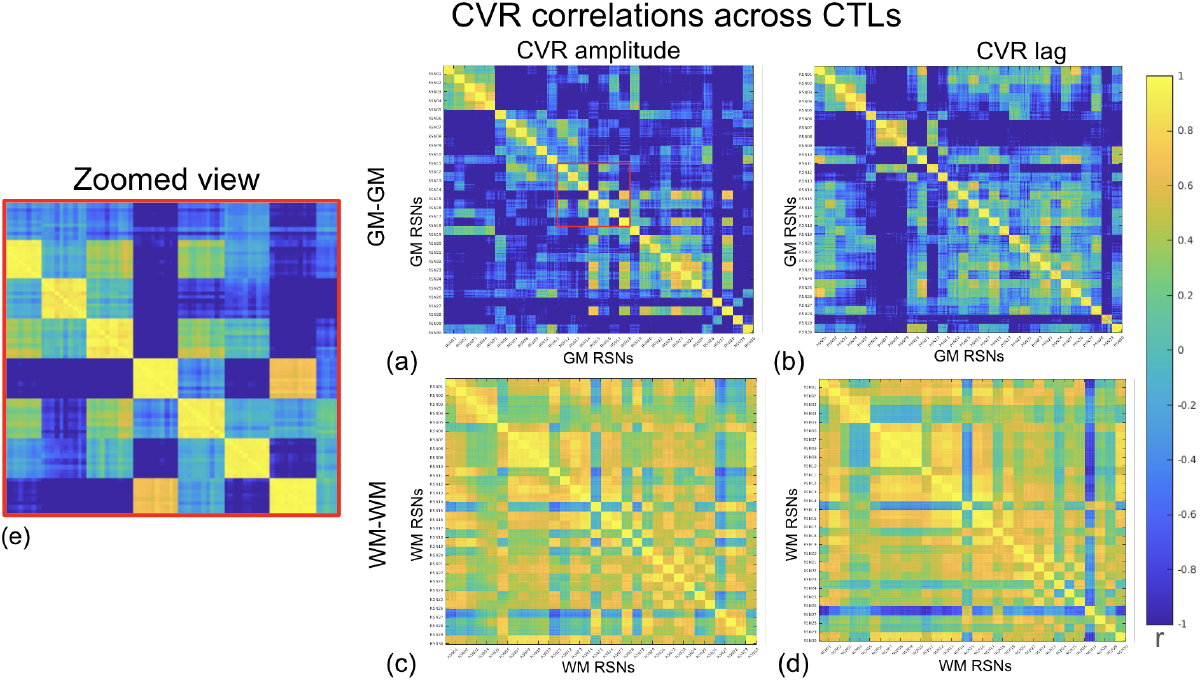
Cross-participant CVR correlation matrix for CTL. (a-b) GM CVR; (c, d) WM CVR. There are 30 grid points per axis, each corresponding to one RSN. Each grid point consists of 10 sub-grid points, representing the 10 randomly subsampled ROIs for each RSN. This is shown in the zoomed view in (e), taken from the red box in (a).

In the whole-brain cluster analysis, the primary observation is that CVR amplitude covariation can be clustered into deep WM (blue), superficial WM (beige), bottom-layer cortex (green), top-layer frontal cortex (purple) and top-layer posterior cortex (red) (**Fig. 8a**). As seen in **Fig. 8b**, the distinguishing features across CVR amplitude clusters are the extent of variation across participants. The corona radiata cluster (shown in green-yellow), for example, is characterized by medium variation across participants, whereas the minor-major forceps cluster (shown in purple) is characterized by large cross-participant variations. The lag-based clusters largely segregate deep WM (red), superficial WM (red) from deep GM (purple) and cortex (blue) (**Fig. 8c**). Unlike the CVR amplitude clusters, the CVR lag clusters are not distinguished by the scale of cross-participant variations, but rather by the patterns of variations (**Fig. 8d**). That is, for instance, while the CVR lag in the purple cluster is greater than the global mean in Participant 2, it is lower than the global mean lag in Participant 2 in the green cluster. Both sets of clusters are symmetrical across hemispheres.

**Figure 8.**
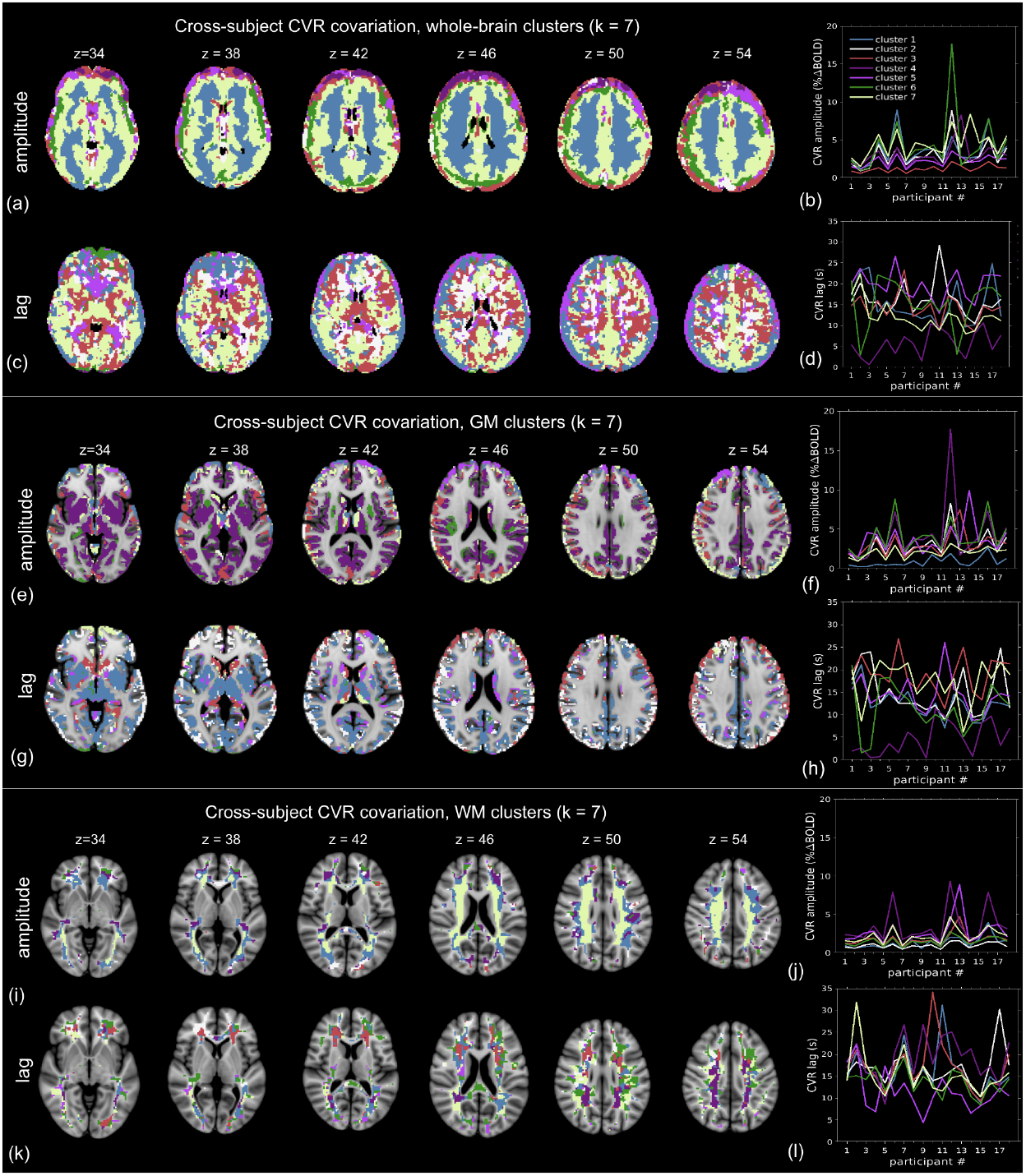
Cross-participant (CTL only) CVR covariation clusters derived based on 7-cluster k-means algorithm. Different clusters are colour-coded for whole-brain (a,b), GM only (c,d) and WM only (e,f). The clustering is based on CVR amplitude patterns (a, c, e) and CVR lag patterns (b, d, f). Note that all values in cluster plots (b, d, f, h, j, l) are based on CVR amplitude or lag. The z value indicates slice coordinates in MNI152 space.

**Figure 9.**
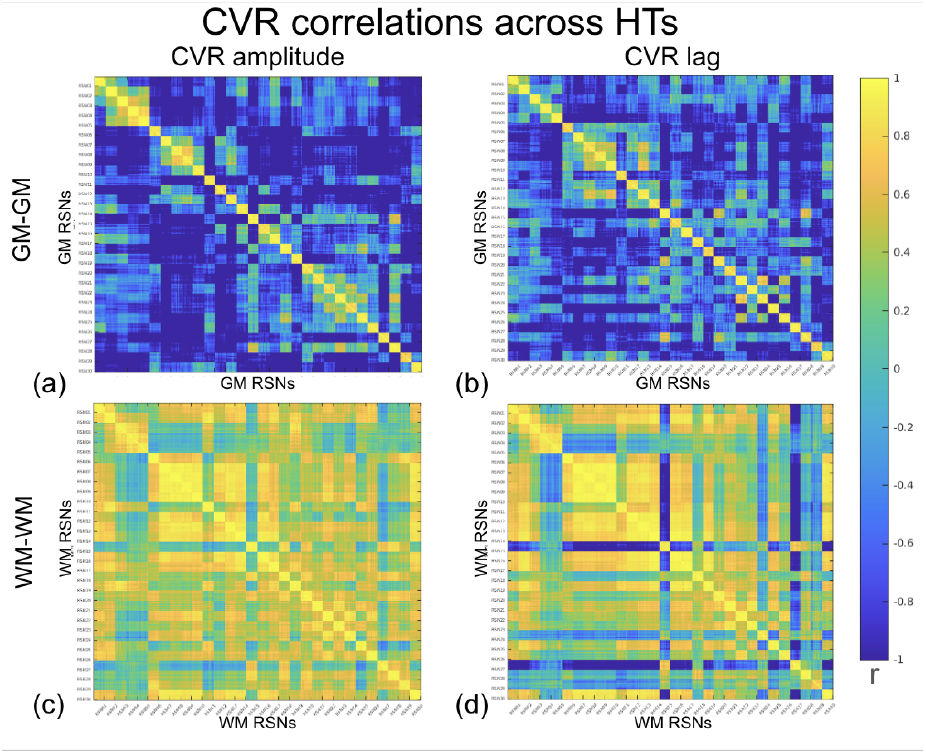
Correlation matrix for HT: GM (top) and WM (top) in the hypertension (HT) group. (a-b) GM CVR; (c, d) WM CVR. There are 30 grid points per axis, each corresponding to one RSN. Each grid point consists of 10 sub-grid points, representing the 10 randomly subsampled ROIs for each RSN.

Focusing on CVR amplitude covariation in GM only, we find a finer pattern that places deep GM and bottom-cortical layer into a single cluster (purple), while the middle-cortical layer (red) and top-cortical layer (beige) continue to be separated (**Fig. 8e**). The thalamus and caudate form a separate cluster from the other deep-GM structures. When k of 17 and 30 are used, the GM is further divided into more layers from superficial to deep, but the general patterns remain consistent (**Fig. S3-4** in Supplementary Materials). Conversely, focusing on CVR amplitude in WM alone, we find a prominent cluster resembling the corona radiata (beige), in addition to clusters comprising the major and minor forceps (blue) (**Fig. 8i**). In the CVR lag cluster maps, these amplitude-based clusters are subdivided into different clusters (**Fig. 8k**). These patterns are consistent in CVR amplitude and lag for both GM and WM when more clusters are used (**Fig. S1-6** in Supplementary Materials). Overall, while differences in the scales of CVR variations across participants dominate the clustering of CVR amplitude, participant-specific CVR lag patterns dominate the CVR lag clustering.

As shown in **Fig. 10**, the CVR lag correlation differences between CTLs and HTs are significant in the correlation between the left middle central (right sensorimotor hand) network and (1) inferior central (sensorimotor hand), (2) superior parietal (dorsal attention), (3) posterior superior parietal, (4) dorsal default-mode, (5) right frontal-temporal parietal 1, (6) right frontal-temporal parietal 3 and (7) cingulo-opercular networks. That is, compared to those of the CTL group, the CVR lag values in HT group exhibit weaker correlations across these networks, while the within-network correlations are preserved.

**Figure 10.**
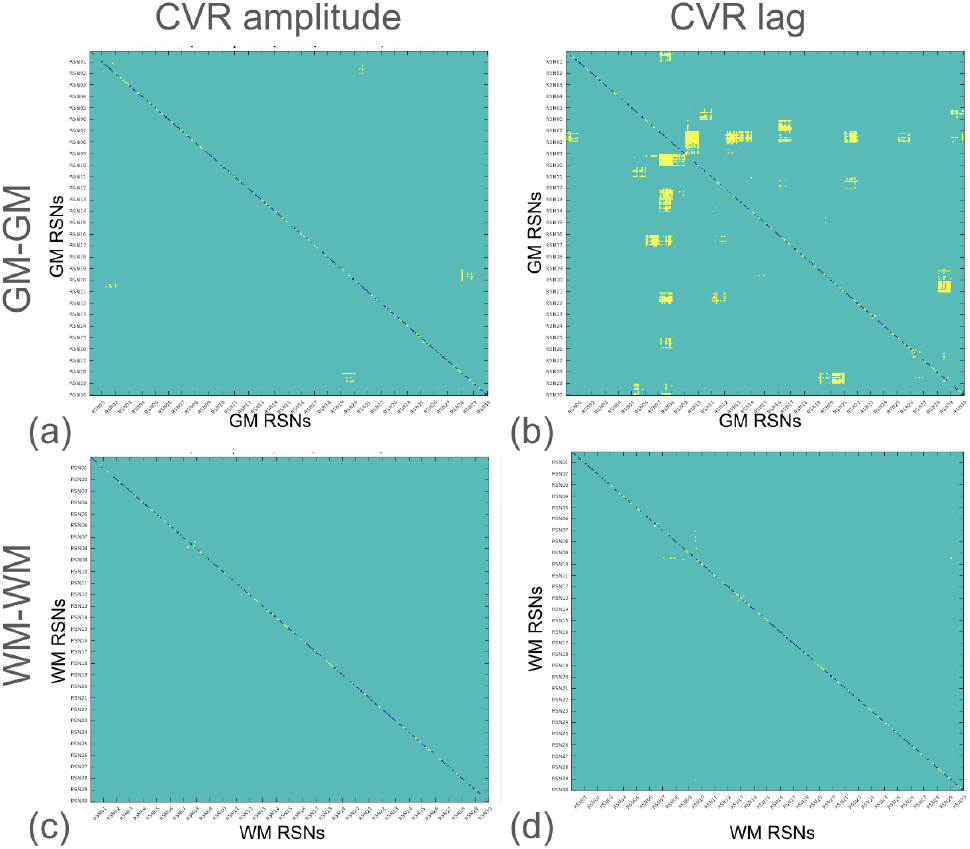
CTL-HT difference matrix. Correlation matrix differences in GM. Each grid point consists of 10 sub-grid points, representing the 10 randomly subsampled ROIs for each RSN. The correlation structures in CVR amplitude are very similar, but the structures in CVR lag display significant differences (p < 0.01), as indicated in yellow.

## Discussion

In this work, we used BH CVR data in healthy older adults and age-matched patients with HT to demonstrate that:

1. The GM brain network structure identified in the Functionnectome coincides with the topography of cross-participant CVR covariations in the GM, while the WM does not display such network structures;
2. Even in the presence of a vascular-modifying pathology such as hypertension, the network nature of cross-participant CVR covariations largely remains unaltered.
3. Both the GM and WM also exhibit different scales of cross-participant CVR amplitude variations depending on tissue layer, as well as CVR lag variations depending on tract (in the WM).

### Regional differences in GM CVR

The work of Tong et al., Bright et al, and Chen et al. laid the foundation for a new way to understand RSNs (Bright et al., 2020; Tong et al., 2015). While these previous works illustrated systemic and vascular-driven correlational spatial structures that are reminiscent of RNS, in this work we tie these spatial structures specifically to the amplitude and timing of CVR. This finding strongly suggests that CVR properties across the brain are segregated by RSNs. Since our results are based on the Funcnectome GM ROIs, the implication is that cortical regions that demonstrate BOLD signal synchrony and are connected by known WM tracts also share CVR properties. In a given participant, certain networks may exhibit higher-than-average or lower-than-average CVR relative to other regions, while in another participant, these trends may be reversed. Thus, the spatial distribution of CVR is not consistent across participants. However, the key takeaway of our findings is that if one node in the RSN of a participant has higher-than-average CVR, so too will the remaining nodes in that RSN. Likewise, if one node in the RSN of a participant has a shorter-than-average CVR lag, so will the remaining nodes in that RSN. The parallel is true for regions demonstrating lower-than-average CVR or longer-than-average CVR lag. Thus, our findings demonstrate that the RSNS tie together the CVR signatures of these spatially distant nodes of RSNs, and each network has a specific CVR signature that distinguishes it from other networks. Possible origins of such network structures will be discussed in a later section.

### Regional differences in WM CVR

In the context of the debated WM fMRI, it is noteworthy that the network structure observed in the GM was not reproduced in the WM. While certain hub regions in the GM are also reflected in the WM correlation matrix, they are but a shadow of their GM counterparts. In fact, the overwhelming trend in the WM is a lack of RSN differentiation. It is possible that the correlations between WM regions are contaminated with noise. After all, the BOLD signal amplitude in the WM is much lower than that of the GM, given the lower CBF (⅓ that of GM) and CVR (¼ that of GM) (van Zijl et al., 1998). It is also possible that WM CVR is better defined by another spatial parcellation. However, another explanation for this finding is that in the WM, CVR amplitude covaries according to depth (**Fig. 8i**) while CVR lag covaries according to tract (**Fig. 8k**). This dependence on tract, coupled with the fact that WM CVR patterns are not well explained by network association, leads us to think that perhaps in the WM, there could be measurement biases in CVR timing due to fibre orientation and the fact that T2* fluctuations are orientation dependent. The WM networks in the Functionnectome are not segregated strictly by tract, as seen in **Fig. 8k**, and this is to be expected, as functional networks are not fully explicable by monosynaptic connections.

These findings seem to run counter to our finding of a significant correlation between CVR amplitude and lag in GM and associated WM ROIs (**Fig. 11**), which suggests a link between GM and WM CVR. The correlations, albeit weak, are significant and found in both CTL and HT participants. A natural question is, if GM and WM CVR are not driven by the same spatial synchronies, then why are they still correlated? These associations, albeit weak, suggest a common vascular supply route shared by GM and WM that is in part governed by their network membership, and should be further explored in a larger data set.

**Figure 11.**
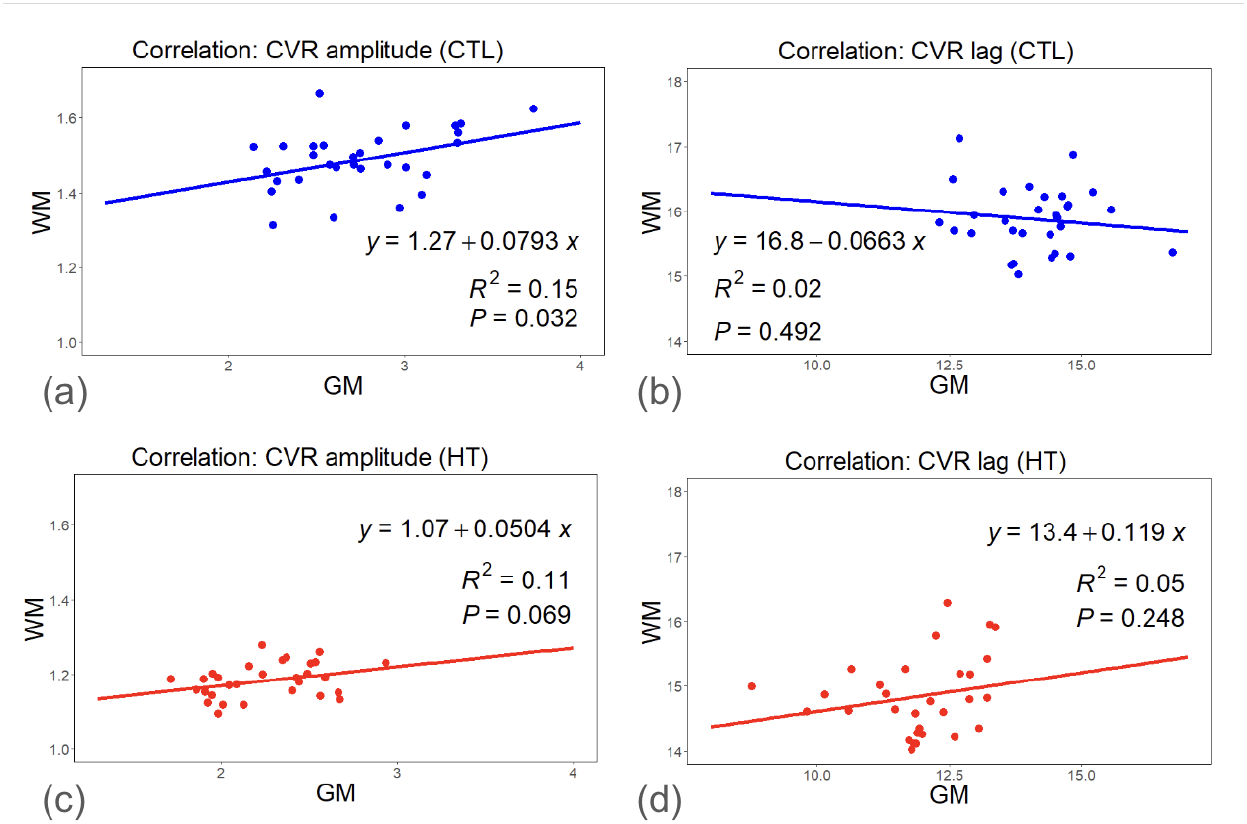
Relationship between GM and WM CVR in CTLs and HTs (scatter plots), for testing the existence of vascular routes that connect the GM and WM. Each dot represents one network ROI in the Functionnectome. Both CVR amplitude and lag are averaged across participants for each ROI. Significant but weak associations are identified between CVR amplitude in the GM and WM networks, in both CTL and HT. There was negligible association between CVR lags in GM and WM networks.

### Hypertension as a model of CVR abnormalities

It is well established that chronic HT leads to an increase in vessel wall stiffness and thickness, reducing the elasticity of blood vessels (Matova and Vihert, 1976). What makes this an interesting test case is that CVR is well known to be altered in patients with HT (Li et al., 2021; Webb and Werring, 2022). In our recent work (Nanayakkara et al., 2023), we demonstrated that HT is associated with a significant reduction in CVR lag, though not in CVR amplitude. This is consistent with the notion of HT leading to increased pulse-wave velocity (Perret and Sloop, 2000).

In our HT group, GM CVR and lag both strongly covary within RSNs, just like in the CTL group (**Fig. 7**). Moreover, there is no significant difference between HT and CTL in terms of the GM and WM CVR amplitude covariation patterns. The only difference in HT compared to the CTL group is the GM CVR lag in HT group exhibits weaker cross-network correlations in these networks, as shown in **Fig. 10**,. All ROIs associated with these differences are consistent with regions exhibiting reduced CVR lag, as reported in our previous work (Nanayakkara et al., 2023, n.d.), and are also linked to the right sensorimotor network. This finding of reduced spatial covariation in CVR in the HT group is also consistent with findings of impaired neurovascular coupling in the frontal and parietal brain regions (Bu et al., 2018). However, given that many of these inter-network covariation patterns are weak (see **Fig. 9b**), and that the M/F ratios are different between the CTL and HT groups, we will not over-interpret the differences between HT and CTL (Vandewouw et al., 2021) groups. Thus, our findings demonstrate that the spatial covariation structure in CVR is maintained in the presence of a vascular pathology that affects CVR. It will certainly be interesting to extend this investigation to other vascular pathologies.

### Possible origins of network structure in CVR properties

Taken together, this paper presents ample evidence to suggest that GM CVR amplitude and lag both covary by functional network. Our results confirm and extend previous findings (Bright et al., 2020; Chen et al., 2020) in that there are indeed networks within the vasoactive system. Moreover, our findings also confirm and extend the findings of Tong et al. (Tong et al., 2015) in that vascular delay can follow a network pattern. A natural follow-up question is why CVR exhibits network properties. The neuronal basis of RSNs has been well established, and there is strong evidence for these RSNs to be sustained by fibre connections (van den Heuvel et al., 2009). Our findings raise many questions regarding the existence and development of RSN-segregated vascular networks. For example, are CVR properties of a given vascular network linked to these neuronal connections? Why is it that each RSN characterized by slightly different CVR properties? The parallel organization of arteries and nerves was first described by Andreas Vesalius (Carmeliet and Tessier-Lavigne, 2005), for which our data may provide experimental evidence.

While neurogenesis proceeds from the WM towards the cortical surface, angiogenesis proceeds from the cortical surface inwards in development (Biswas et al., 2020). Endothelial tip cells (Wälchli et al., 2015), a specialized subtype of endothelial cells that guide the leading edge of angiogenesis, guide the blood vessel sprout toward gradients of angiogenic growth factors such as vascular endothelial growth factor (VEGF) or basic fibroblast growth factor (Carmeliet and Tessier-Lavigne, 2005). Tip cells are followed by endothelial stalk cells, which proliferate and form the vascular lumen, thereby elongating the vascular sprout (Carmeliet and Tessier-Lavigne, 2005). The release of VEGF, in turn, is modulated by the amount of neuronal/cellular metabolism (Zafar et al., 2017), and the endothelial stalk cells form the endothelium where CVR takes place (Mazurek et al., 2017).

CVR is mainly mediated by calcium (Ca^2+^) signalling the vascular smooth muscles and astrocytes, pericytes as well as by the blood-brain barrier (BBB). During a vascular challenge, smooth muscles relax to allow the blood vessels to widen, increasing blood flow to the activated area (Bank et al., 1995). Smooth muscle cells also release signaling molecules like nitric oxide, which further promote vasodilation and blood flow (Levine et al., 2012). A compromised BBB can lead to increased vascular permeability, potentially disrupting blood flow regulation and hindering vasodilation during CVR (Bernardo-Castro et al., 2020; Price et al., 2016). Thus, it is conceivable that properties of the BBB can modulate CVR amplitude and delay. Studies of the mouse model demonstrate that BBB maturation does not complete until after synaptogenesis and neuronal-activity dependent synaptic pruning begin (Biswas et al., 2020). Interestingly, tight junctions, protein complexes that form the backbone of the BBB, might vary slightly between brain regions (Hupe et al., 2017). Moreover, endothelial cells, which line the blood vessels and form the BBB, might exhibit some diversity in gene expression patterns across brain regions (Geudens and Gerhardt, 2011). This could potentially lead to functional variations and spatial differentiation in the BBB. Additionally, pericytes, which wrap around endothelial cells and maintain the BBB, respond to signaling by endothelin-1 (ET-1) and nitric oxide (NO), leading to vasoconstriction and vasodilation, respectively. Studies suggest that pericytes can vary in density, morphology and marker expression across the brain, such as between the cortex and the deep GM (Yamazaki and Mukouyama, 2018). Finally, astrocytes communicate with both smooth muscle cells and endothelial cells (cells lining the BBB) using various signaling molecules. These signals can influence blood flow regulation and smooth muscle contractility during a CO_2_ stimulus (Turovsky et al., 2016).

Despite the spatial heterogeneity, there is currently no clear evidence that the endothelial cells and the BBB differ based on RSNs. However, there is growing evidence that astrocytes exhibit some differences across RSNs. Astrogenesis begins later than both neurogenesis and angiogenesis (Biswas et al., 2020; Bond et al., 2020). Astrogenesis takes place as the transcription of angiogenic markers start to decline (Bayraktar et al., 2020), and marks the transition from angiogenesis to vascular maturation and BBB development. While astrocytes were traditionally viewed as a uniform cell population, recent research suggests they are more diverse than previously thought. Studies suggest that there might be differences in astrocyte morphology and gene expression patterns between brain regions associated with different RSNs (Endo et al., 2022). Astrocytes may release specific signaling molecules that influence neuronal activity within a particular network. Furthermore, astrocytic calcium signaling patterns might differ across RSNs, potentially reflecting their role in network-specific functions. A recent review (Khakh and Sofroniew, 2015) highlights that astrocytes display diverse intracellular Ca^2+^ signals that are spatially and temporally unrelated to those in the somata and are distinct within circuits (Srinivasan et al., 2015). The brain-circuit differentiation in astrocyte signalling may be enough to induce RSN-specific CVR signatures.

In the WM, unlike in the GM, the network structure is not apparent, as revealed by the correlation matrices. If the astrocytes in the GM are the main carrier of network specificity, what about astrocytes in the WM? The WM is populated mainly by fibrous astrocytes, which have a more elongated morphology compared to protoplasmic astrocytes and are thought to be specialized for functions related to myelination and supporting WM structure (Lundgaard et al., 2014). Fibrous astrocytes release signalling molecules that influence oligodendrocyte development (Li et al., 2016). In the fetal stage, axonal development precedes neuronal development in the cortex, so presumably, vascular development in the WM precedes that in the GM. However, oligodendrocyte differentiation begins after neuronal differentiation from radial glial cells (Martynoga et al., 2012; Rowitch and Kriegstein, 2010), and there is currently limited evidence to support the differentiation of fibrous astrocytes or oligodendrocytes across brain circuits. Furthermore, it is not completely clear how CVR is mediated in the WM, or if the neurovascular unit in the WM resembles that in the GM (Bernier et al., 2021). Thus, the CVR structure in the WM has henceforth been more difficult to understand.

### Possible origins of tissue-layer structure in CVR properties

Integrating the clustering results with the correlation matrices, it is also interesting to observe that beneath the commonality of network structure, there remain different patterns of spatial covariation in CVR amplitude for different tissue layers, whereas the spatial covariation in CVR lag is more heterogeneous. In the GM, CVR lag covaries according primarily to cortical depth. WM CVR amplitude also follows a layer-like structure, whereas CVR lag a tractwise pattern. Although GM and WM T2* variations have been reported to be orientation dependent (Doucette et al., 2019; Lee et al., 2011; Rudko et al., 2014), here we do not observe GM or WM clustering that could be explained by cortical or fibre orientation. This will be further discussed in the next section.

Does this layer-like structure in CVR revealed in the cluster analysis run counter to the network-based structure revealed in the correlation analyses? While Pearson’s correlation is maximized with high signal covariance and low variance, k-means clustering identifies clusters by minimizing the Euclidian distance between data series. In our case, for two voxels to cluster together, their cross-participant CVR variations may not be correlated, but the scales of these variations must be similar. Thus, in the cluster analysis, we cluster together regions that collectively exhibit large (or small) variations across participants.

With this in mind, the layer-specific clusters can be interpreted as: with each layer, the expected inter-participant variation in CVR differs. As shown in **Fig. 8a and b**, the deep WM layers exhibit greater levels of CVR variations than superficial cortical layers across participants. We know that bottom cortical layers and deep WM layers will be more distal to the arterial blood supply. This finding raises the possibility that CVR amplitude is most variable in poorly perfused regions, and leads us to wonder whether vascular resiliency in the presence of disease is greater in the better perfused brain regions. Furthermore, the aforementioned difference in pericyte properties between cortical and deep GM regions is also consistent with the observed CVR amplitude cluster patterns. CVR lag, on the other hand, appears to reveal vascular delivery routes. This is particularly the case in the WM, due to the tract-like spatial pattern of the clusters. It is interesting to appreciate how the GM CVR lag segregates by “anterior”, “posterior”, “deep” and “superficial”, not seeming to correspond to the lag patterns in WM CVR (**Fig. 8 e-l**).

### Limitations and future work

This work is based on the BOLD contrast mechanism, which provides an indirect measure of CVR. Although BOLD-based CVR is robust and clinically recognized (Chen and Fierstra, 2021; Duffin et al., 2021; Pillai and Mikulis, 2015), we will extend this work to perfusion-based CVR (Halani et al., 2015) in our future work. Furthermore, this work is based on a modest sample of older adults, and thus there could be underlying un-diagnosed subclinical HT in the CTLs. While our findings are clear and robust, we aim to reproduce this analysis in a larger sample of younger adults. Lastly, we hope to confirm these findings in data obtained using passive gas challenges instead of breathholding.

## Conclusions

The overall message of this work is that while CVR may synchronize across distant brain regions that belong to the same functional network, it is also modulated by certain local tissue properties related to tissue-layer/tract membership. This work demonstrates the use of cross-participant covariation patterns in CVR as an informative way of mapping the vascular routes in the GM and WM, and also raises questions about the nature of the widely cited RSNs.

## Acknowledgments

We are grateful to the Canadian Institutes of Health Research (CIHR) [grant number FDN 148398] for financial support (NN, JJC).

## Author contributions

NDN contributed to conceptualisation, data curation, formal analysis, investigation, methodology, software, validation, visualisation, writing—original draft.

LAM and NDA contributed to data curation, writing—review & editing.

JJC contributed to conceptualisation, resources, funding acquisition, data curation, methodology, supervision, writing—original draft, writing—review & editing.

## Data availability

The data that support the findings of this study may be made available only according to terms in the research ethics approval upon request to the corresponding author. The data are not publicly available due to privacy or ethical restrictions.

## Computer code availability

Code will be made available upon request to the corresponding author.

