## Supplementary Materials for "Inter-individual differences in cerebrovascular reactivity are synchronized within functional networks and tissue layers: evidence from healthy older adults and patients with hypertension"

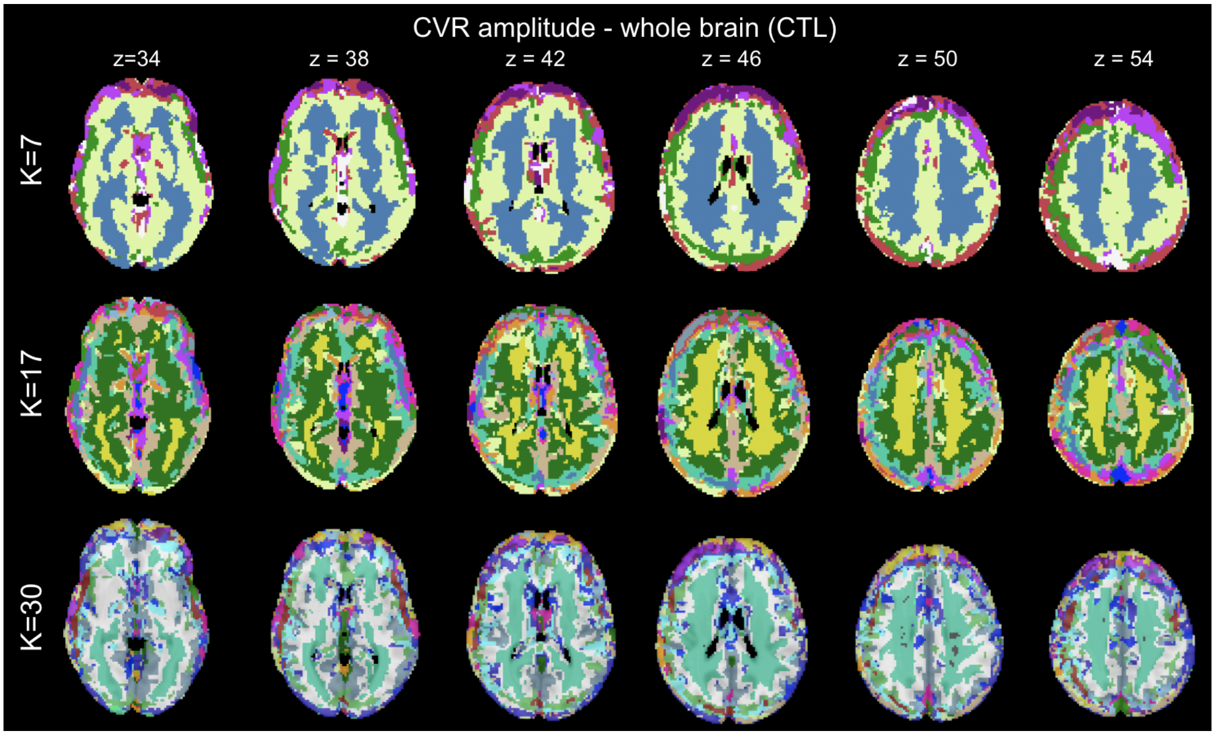


**Figure S1**. **Cross-participant (CTL only) CVR amplitude covariation clusters derived for the GM based on 7, 17 and 30-cluster k-means algorithm.** Different clusters are colour-coded in each case.

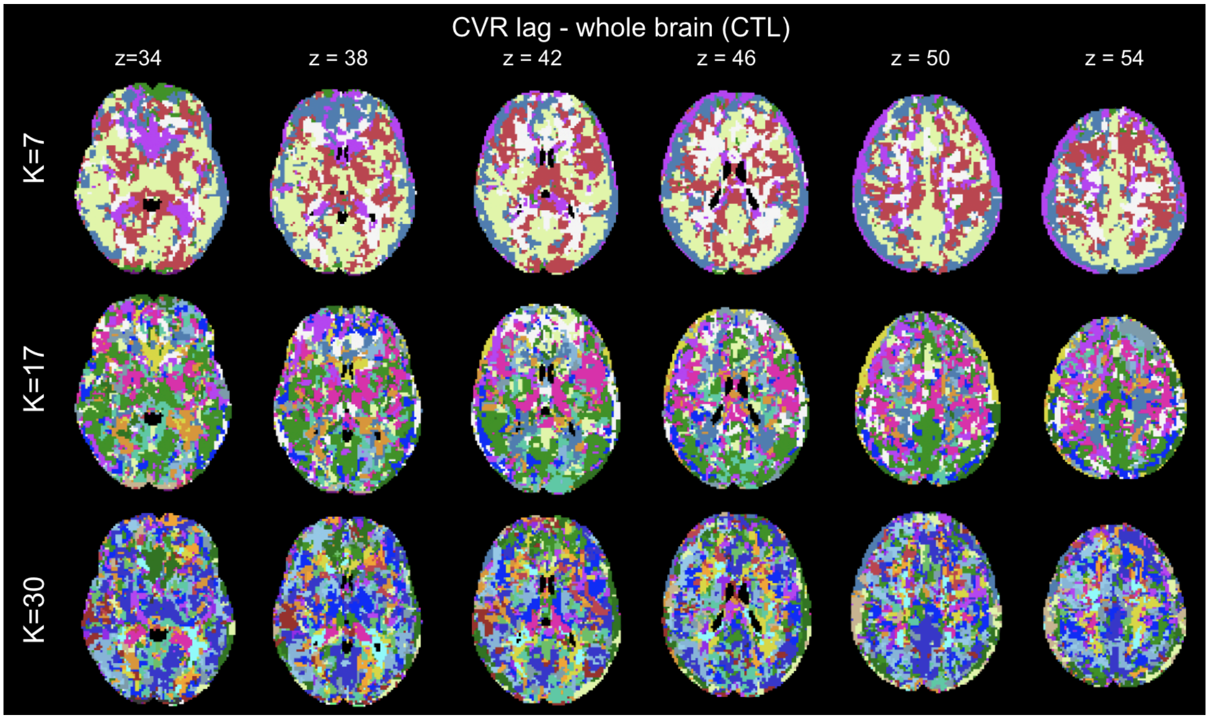


**Figure S2**. **Cross-participant (CTL only) CVR lag covariation clusters derived for the GM based on 7, 17 and 30-cluster k-means algorithm.** Different clusters are colour-coded in each case.

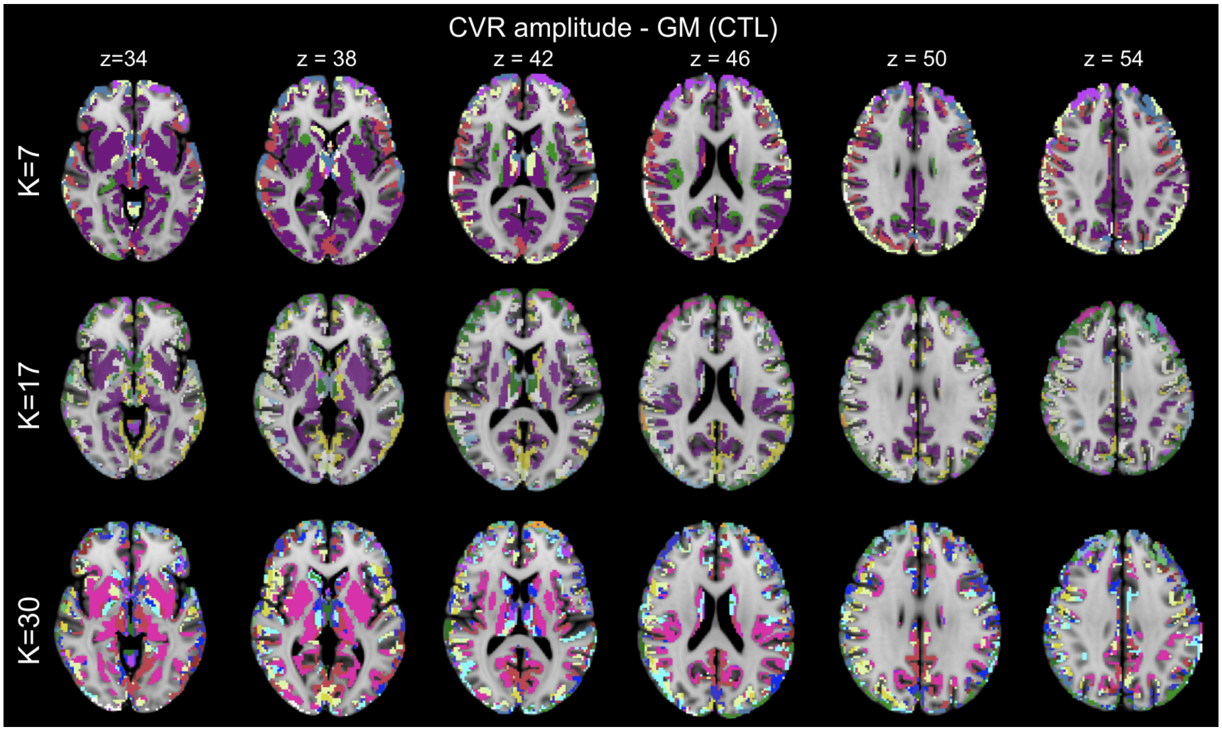


**Figure S3**. **Cross-participant (CTL only) CVR amplitude covariation clusters derived for the GM based on 7, 17 and 30-cluster k-means algorithm.** Different clusters are colour-coded in each case.

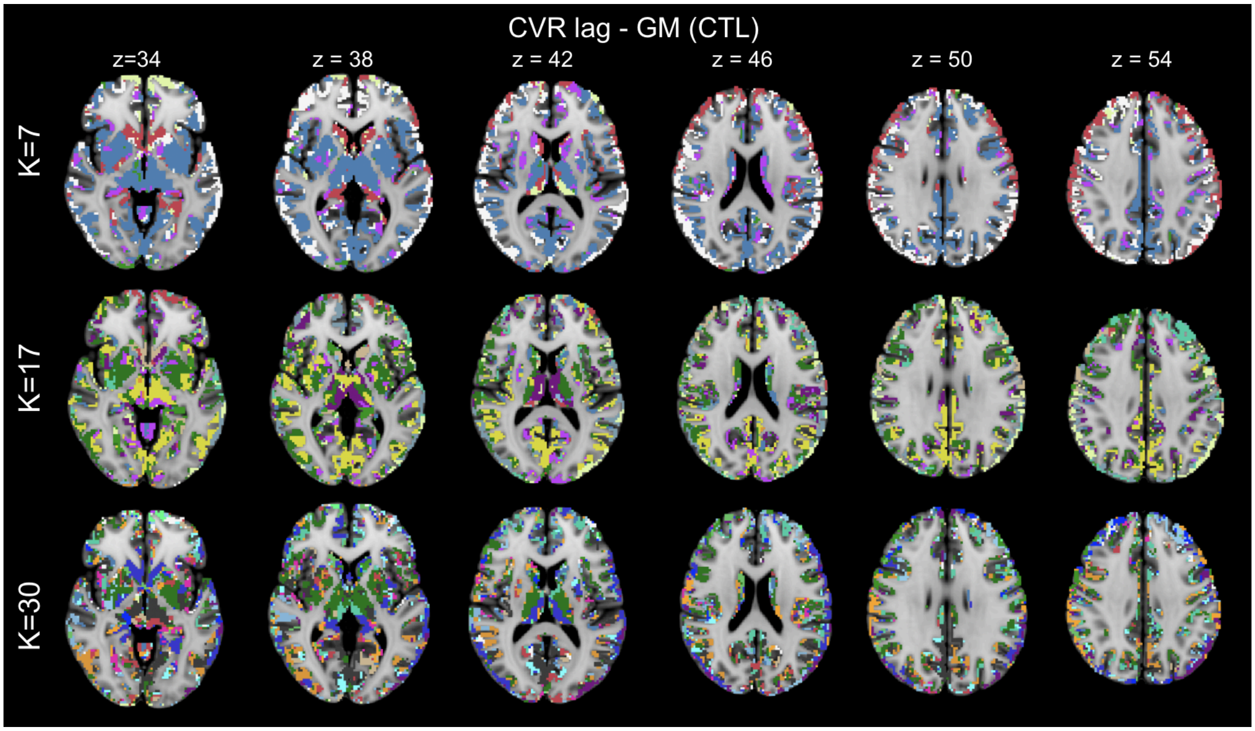


**Figure S4**. **Cross-participant (CTL only) CVR lag covariation clusters derived for the GM based on 7, 17 and 30-cluster k-means algorithm.** Different clusters are colour-coded in each case.

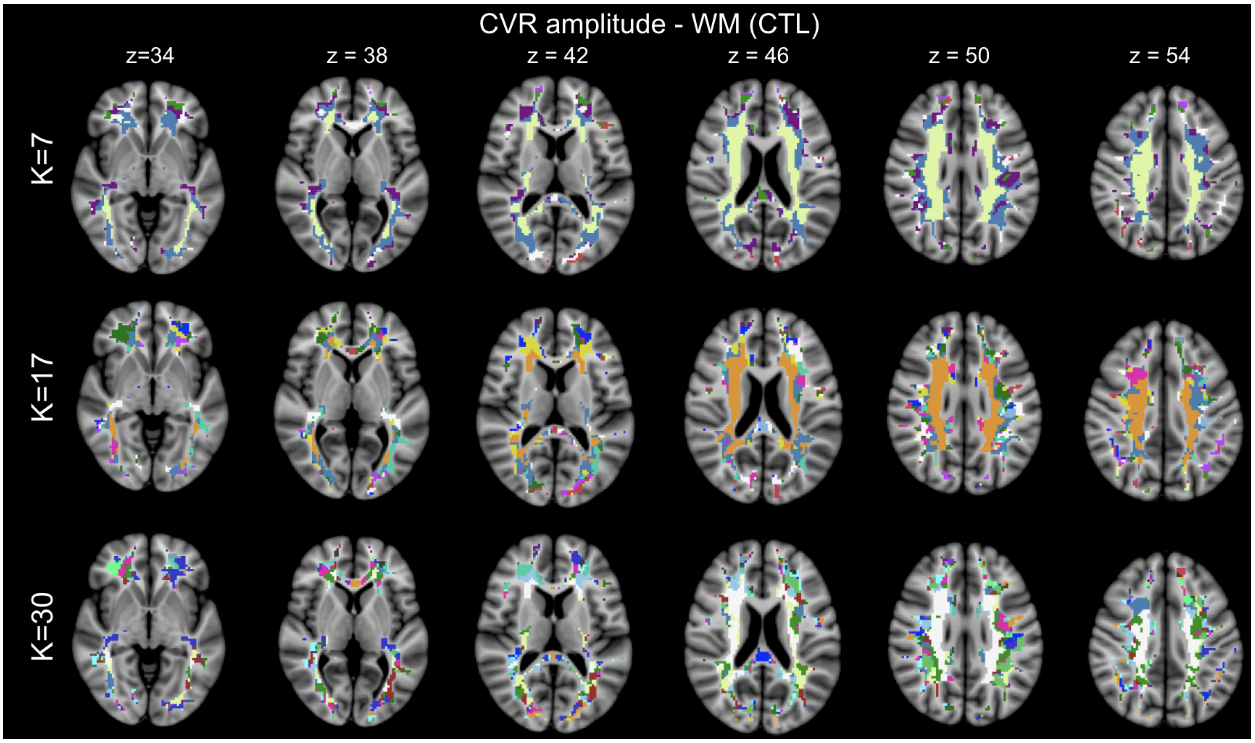


**Figure S5. Cross-participant (CTL only) CVR amplitude covariation clusters derived for the WM based on 7, 17 and 30-cluster k-means algorithm.** Different clusters are colour-coded in each case.

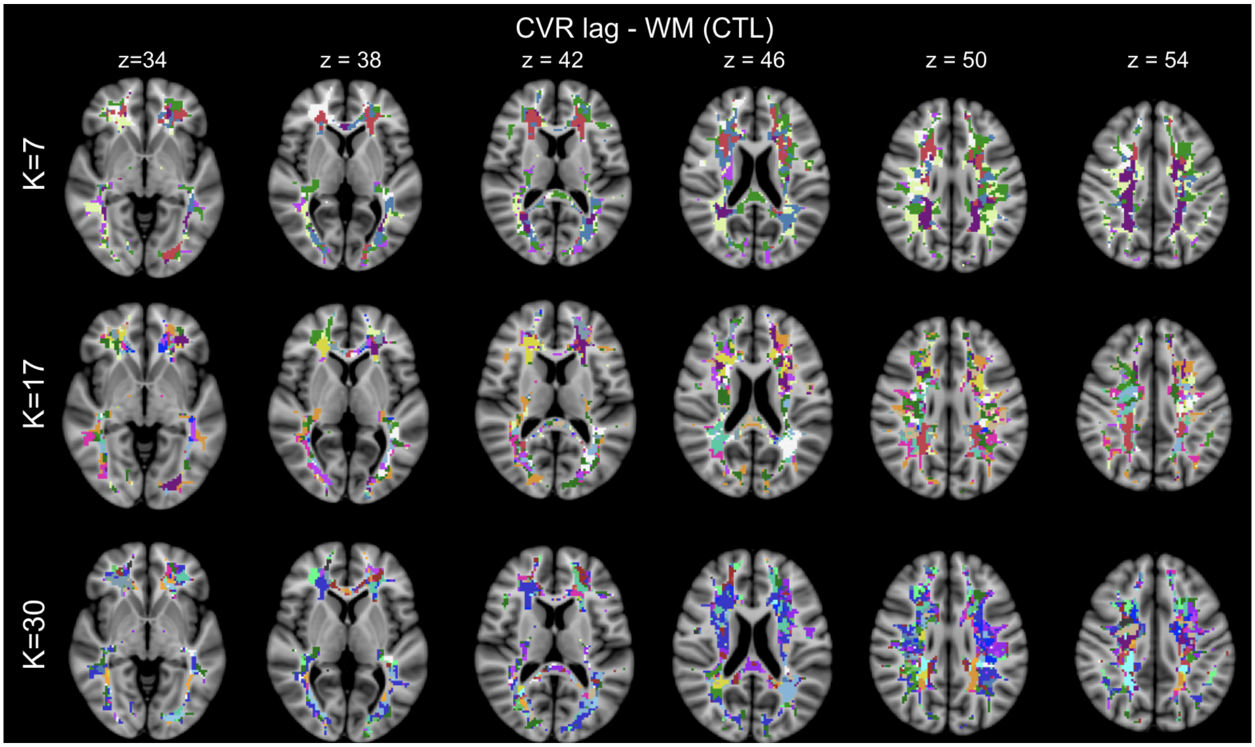


**Figure S6. Cross-participant (CTL only) CVR lag covariation clusters derived for the WM based on 7, 17 and 30-cluster k-means algorithm.** Different clusters are colour-coded in each case.
